# PAT: An Image Analysis Tool for Automated Scoring of Pollen in Alexander-Stained Anthers

**DOI:** 10.64898/2026.05.07.723495

**Authors:** Darya Volkava, Vivek Kumar Raxwal, Karel Riha

## Abstract

Quantitative pollen viability analysis is a critical but labor-intensive step in plant reproductive biology. Existing deep-learning Segment Anything Models (SAM) fail to reliably segment viable pollen in Alexander-stained anthers. To address this, we fine-tuned an existing Cellpose-SAM model for pollen segmentation. We integrated it into PAT (Pollen Analysis Tool), a cross-platform desktop application. PAT features instance segmentation with interactive quality control, an in-app model retraining module, and publication-ready statistical outputs. We deployed PAT in an EMS suppressor screen of semi-sterile Arabidopsis *smg7-6* mutants, enabling efficient candidate prioritization for whole genome sequencing and mapping candidate mutation. This screen led to the identification of a point mutation in CAP-D2 (*capd2-2*), a Condensin I subunit, that rescues the *smg7-6* meiotic phenotype. Notably, mutation in a Condensin II subunits (CAP-D3 and CAP-H2) does not confer rescue. Further characterization suggests the *capd2-2* allele is hypomorphic, showing no defects in vegetative growth, chromocenter compaction, or transposable element silencing. Collectively, we demonstrate that accessible AI tools have the potential to bridge gaps in plant phenotyping and accelerate the pace of biological discovery.

**Highlight:** We combined AI-powered image analysis with an easy-to-use desktop app to automate plant pollen counting, then used it to identify a new genetic suppressor of meiotic defects.

## Introduction

Pollen viability is a key determinant of male reproductive success in flowering plants. It directly influences fertilization efficiency, seed set, and final crop yield. The development and function of the male gametophyte (pollen) is a complex, tightly regulated process that is highly sensitive to both genetic and environmental factors (Borg *et al*., 2009; Marchant and Walbot, 2022). Abiotic stresses such as high temperature, drought, and cold can severely impair microsporogenesis and pollen maturation, often resulting in reduced viability or complete sterility (Hedhly *et al*., 2009; Zinn *et al*., 2010; De Storme and Geelen, 2014; Rieu *et al*., 2017; Qian *et al*., 2025). Genetic mutations disrupting meiosis, or post-meiotic development produce similar effects, frequently causing male sterility (Borg *et al*., 2009; Williams and Mazer, 2016; Marchant and Walbot, 2022). Owing to this sensitivity, pollen viability assessment serves as a practical and reliable tool for studying plant response to stress, screening crop germplasm for abiotic stress tolerance, and for identifying genes involved in pollen development (Prasad *et al*., 2006; Burke and Chen, 2015; Djanaguiraman *et al*., 2018; Ingole *et al*., 2025; Miller *et al*., 2026). Accurate analysis of pollen count and viability is therefore essential for modern plant breeding programs, hybrid seed production, and advancing our understanding of reproductive biology.

Several methods are commonly used to assess the functional and physiological integrity of pollen. Traditional cytological staining techniques, such as Alexander’s stain, tetrazolium salts (TTC or MTT), and fluorescein diacetate with propidium iodide (FDA/PI), use chemical dyes to evaluate membrane integrity, metabolic activity, or enzyme function (Alexander, 1969; Heslop-Harrison and Heslop-Harrison, 1970; Jones and Senft, 1985; Rodriguez-Riano and Dafni, 2000; Tang *et al*., 2025). In vitro germination assays provide a more direct functional test by measuring the pollen’s ability to rehydrate, germinate, and successfully produce pollen tubes in optimized media (Brewbaker and Kwack, 1963; Dafni and Firmage, 2000; Dickinson *et al*., 2018). In contrast, impedance flow cytometry (IFC) offers a modern, label-free, high-throughput approach that analyzes the biophysical and electrical properties of thousands of individual pollen grains per second, delivering rapid and quantitative data on pollen quality (David *et al*., 2012; Impe *et al*., 2019). Nevertheless, Alexander’s stain remains the standard first-line method for routine pollen viability assessment (Crismani *et al*., 2013; Khouider *et al*., 2021; Feng *et al*., 2023; Seear *et al*., 2025). Its continued use stems from its procedural simplicity, high reproducibility, negligible cost, and minimal equipment requirements — only a standard bright-field microscope is needed. This histochemical protocol leverages the differential affinities of acid fuchsin and malachite green to resolve cytoplasmic integrity. Under bright-field microscopy, viable grains sequester acid fuchsin within the intact, metabolically active cytoplasm, yielding a vivid magenta or purple-red coloration. Conversely, aborted or dead grains—devoid of functional cytoplasm—fail to retain the red dye, allowing malachite green to stain the non-viable pollen and debris, a distinct blue-green. This sharp chromatic contrast facilitates the rapid quantification of male fertility across large sample sizes. Because the resulting viability data correlate strongly with physiological germination rates, this method remains an indispensable, high-throughput proxy for evaluating male reproductive fitness (Atlagić *et al*., 2012; Devasirvatham *et al*., 2012; Stokes and Geitmann, 2025).

Despite the utility of Alexander’s stain for rapid fertility assessments, the transition from qualitative histological observation to high-throughput quantitative phenotyping remains technically challenging. Current methodologies largely rely on manual visual scoring, which introduces significant inter-observer bias. It also restricts the number of samples that can be produced, hence limiting the statistical power of the obtained data. This constraint is particularly evident in large-scale genetic studies, such as genetic screens, genome-wide association studies, or studies assessing the effect of the environment on plant reproduction (De Muyt *et al*., 2009; Tsuchimatsu *et al*., 2020; Capitao *et al*., 2021; Guo *et al*., 2023; Crhak Khaitova *et al*., 2024; Kasthurirengan *et al*., 2025). Therefore, specialized automation is required to replace manual counting with a more accurate, reproducible and statistically robust approach.

To overcome these challenges, we developed the Pollen Analysis Tool (PAT), an open-source platform designed for the objective, high-throughput quantification of pollen viability from Alexander-stained samples. PAT leverages a customized Cellpose-Segment Anything Model (CPSAM) architecture (Marks *et al*., 2025), which we optimized through training on a curated dataset of diverse Alexander-stained images to ensure robust segmentation across varying optical qualities. The application integrates an intuitive graphical interface with an interactive quality-control suite and a ‘human-in-the-loop’ refinement module. We validated the utility of PAT by conducting a forward genetic screen of an ethyl methanesulfonate (EMS)-mutagenized Arabidopsis population. Specifically, we employed PAT to identify suppressors of the male infertility phenotype, which stems from aberrant meiotic exit and subsequent mapping of the causative suppressor locus, demonstrating PAT’s efficacy in large-scale genetic discovery.

## Results

### 3.1 The cellpose CPSAM model shows density-dependent errors on Alexander-stained anther images

To enable high-throughput pollen viability quantification, we first evaluated whether existing Cellpose segmentation model (CPSAM) could directly process Alexander-stained *Arabidopsis thaliana* anther images. CPSAM is the current state-of-the-art generalist model for microscopy segmentation (Stringer *et al*., 2021; Marks *et al*., 2025). Given CPSAM robust gradient-flow architecture and its established ability to handle diverse cell morphologies with minimal parameter tuning (Stringer *et al*., 2021; Marks *et al*., 2025), we considered it a strong a priori candidate for high-throughput pollen analysis.

When we tested CPSAM across images spanning the full range of pollen densities—from densely packed fertile anthers to near-empty sterile ones—the model exhibited systematic, density-dependent errors (**Fig. 1**). In dense, fertile anthers, it consistently under-counted grains and in sparse, low-viability anthers, it significantly over-counted by failing to distinguish between viable pollen and confounding elements like non-pollen somatic cells. We attempted to correct these errors by adjusting the model’s diameter and flow threshold parameters; however, this produced inconsistent improvements (**Fig. 1**). Optimal settings varied substantially across staining batches and genotypes, requiring time-consuming manual inspection for each image. For datasets involving hundreds of images, such manual tuning offered no practical advantage over traditional manual counting.

**Fig. 1.**
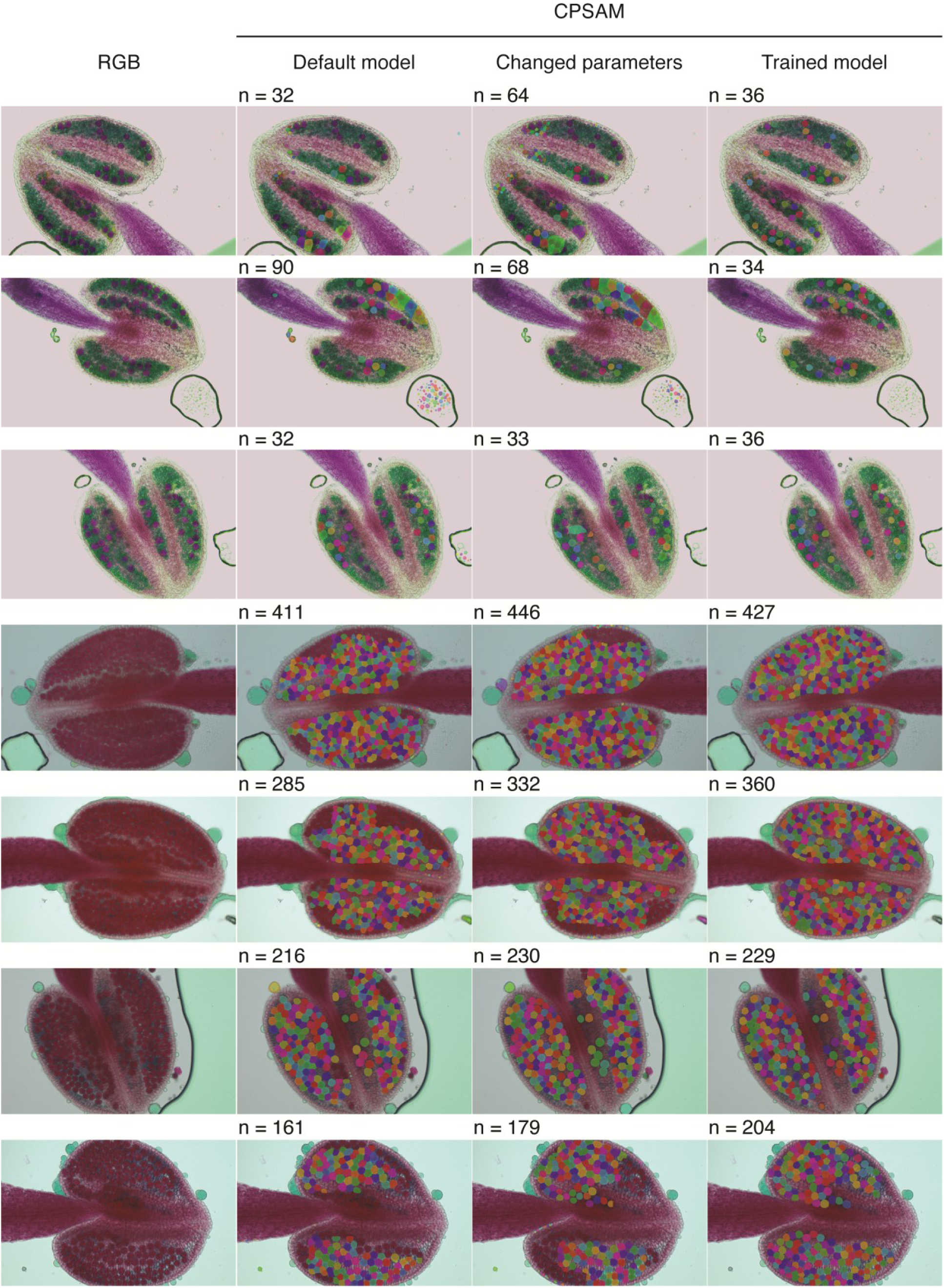
Cellpose performance on Alexander-stained anther cross-sections across varying pollen densities. Representative cross-sections of Alexander-stained anthers showing a range of pollen densities, from low (top rows, light staining) to high (bottom rows, dense red staining). Each row presents the same anther image processed under four conditions (left to right): (i) original unprocessed image, (ii) segmentation using the Cellpose default CPSAM model, (iii) segmentation using the default model with manually adjusted parameters, and (iv) segmentation using a custom-trained Cellpose model. Colored overlays represent individually segmented pollen grains, with each color denoting a unique detected instance; the number of detected objects (n) is indicated above each segmented panel. The default model (columns 2–3) produces spurious detections and fails to reliably segment pollen grains even after parameter tuning, particularly in high-density samples. The custom-trained model (column 4) consistently detects pollen across all density conditions while eliminating false positives

These limitations established a clear need for a domain-adapted model capable of specifically identifying viable pollen. We therefore curated a dedicated training dataset of 65 Alexander-stained anther images encompassing more than 20,000 manually annotated pollen grains. This dataset reflects the diversity of conditions encountered in high-throughput screens, including various densities and staining qualities. We annotated all images with per-grain instance segmentation masks and trained the CPSAM model for 200 epochs. The resulting custom-trained model successfully learned to differentiate the distinct morphological features of viable pollen from other purple-stained structures, achieving high accuracy across both high- and low-density samples (**Fig. 1**).

### 3.2 Quantitative validation of the custom-trained model justifies its use for automated analysis

To determine if a custom-trained model could provide the accuracy necessary for high throughput tasks, we compared its performance against manual “ground-truth”. We used a validation set of manually curated 50 images that captured the full range of biological variation seen in our experiments, from sterile mutants with zero pollen to high-density wild-type anthers with over 300 grains. This validation set was used to compare and benchmark default CPSAM and custom trained CPSAM.

The custom-trained model performed almost identically to the human expert across all samples (**Fig. 2A**). In contrast, the standard CPSAM model performed unreliable when processing the complex textures of Alexander-stained anthers. It consistently over-counted in sterile samples by mistaking somatic debris for pollen and severely under-counted in fertile samples by failing to separate tightly packed grains. When we quantified these errors, the custom model was highly accurate: approximately 90% of its counts were within difference of just 5 grains of the manual count and 95% were within 10 grains (**Fig. 2B**). CPSAM, however, exceeded 30-grain margin of error in more than 60% of images, making it unsuitable for deploying it for automatic pipeline (**Fig. 2B**). Next, we plotted the residual error distribution, which represents the raw difference between the manual ‘ground-truth’ and the model’s prediction (**Fig. 2C**). The custom model exhibited virtually no bias and model count difference from manual count centered around zero. In contrast, CPSAM showed a much wider and unpredictable range of errors, in some cases deviating from the true count by more than 100 grains.

**Fig. 2.**
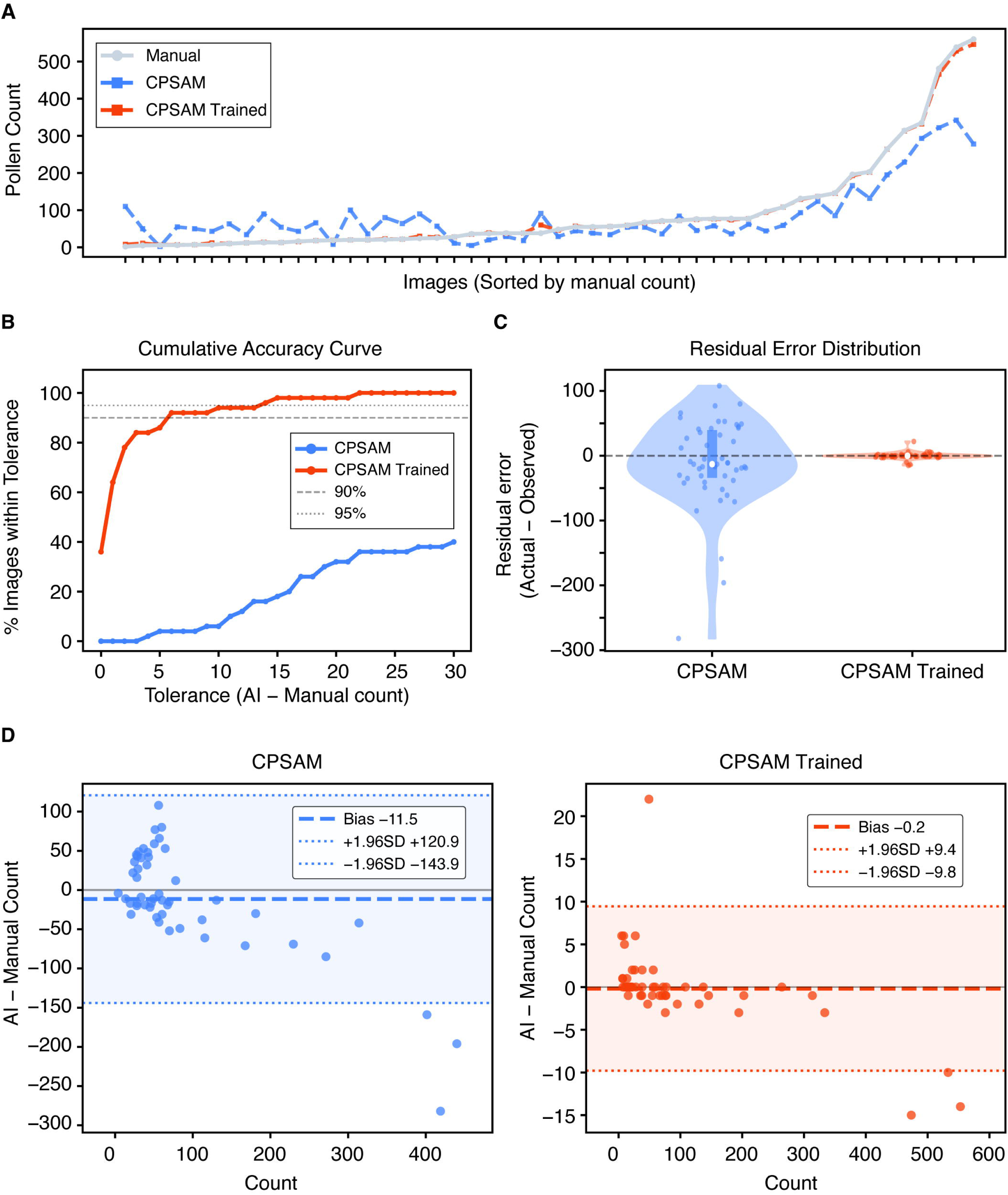
Quantitative comparison of CPSAM and the custom-trained model against manual pollen counts. **(A)** A line plot showing pollen counts per image in ascending order from manual count (light grey), the custom model (orange), and default CPSAM (blue). **(B)** Cumulative accuracy curves showing the percentage of images within a given error tolerance (grains). The custom model reaches 95% accuracy within a 10-grain tolerance. **(C)** Violin plots of residual errors (manual – AI count). The custom model exhibits negligible bias compared to the broad, asymmetric error distribution of CPSAM. **(D)** Bland–Altman plots showing mean bias (dashed line) and 95% limits of agreement (LoA; dotted lines). The custom model narrows the LoA more than tenfold (−9.8 to +9.4 grains) compared to CPSAM (−143.9.0 to +120.9 grains), demonstrating high agreement with manual count.

To determine if the model could reliably replace manual scoring in a large biological context, we performed Bland–Altman analysis (Bland and Altman, 1986) to evaluate the limits of agreement between methods (**Fig. 2D**). In this context, the limits of agreement define the range within which 95% of the differences between manual and automated counts are expected to fall. While CPSAM exhibited unacceptably broad limits (−143.9.0 to +120.9 grains), our fine-tuned model narrowed this range more than tenfold (−9.8 to +9.4 grains) (**Fig. 2D**). These results demonstrate that while general-purpose AI models struggle with the overlapping signals of pollen and anther tissue, our custom training reduces counting errors to a negligible level. This high degree of precision established the necessary foundation to develop a dedicated, automated analysis tool.

### 3.3 PAT: an integrated graphical desktop tool for pollen analysis

High numerical performance alone does not guarantee broad adoption. While deep-learning frameworks like Cellpose offer state-of-the-art accuracy and provide excellent documentation for installation, their routine use in a biological laboratory often requires a level of computational expertise that can be a barrier to entry. Implementing command-line workflows, managing Python environments, and handling complex software dependencies across different operating systems can be daunting. To complement the power of deep learning model with a more accessible interface, we developed PAT (Pollen Analysis Tool), a graphical desktop application implemented in Python using the PyQt6 framework (**Fig. 3**). PAT simplifies the deployment of these advanced model by providing a native, “one-click” environment on Windows, macOS, and Linux that requires no programming knowledge.

**Fig. 3.**
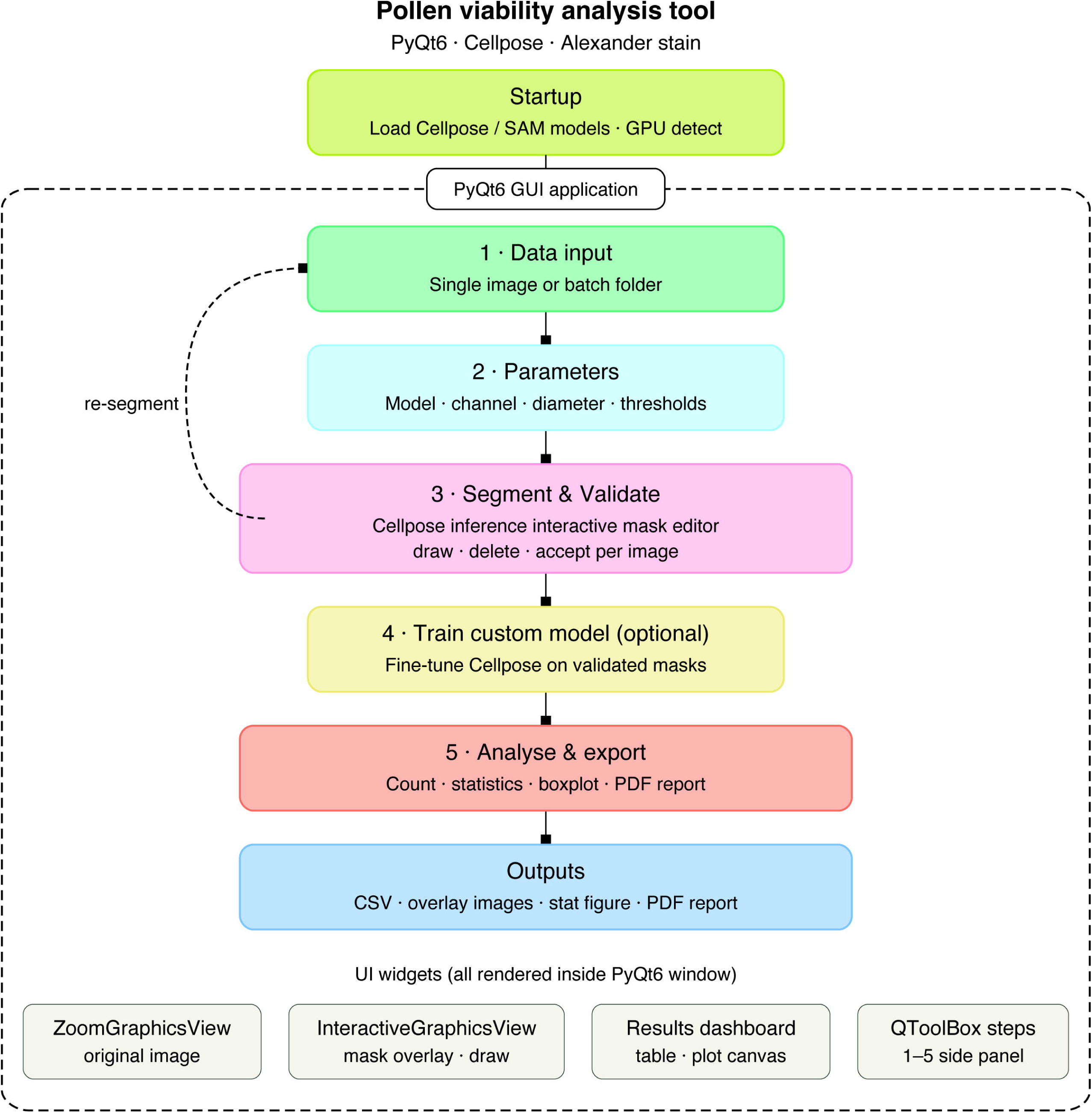
Architecture and workflow of the Pollen viability Analysis Tool (PAT). Schematic overview of PAT, a PyQt6-based desktop application integrating Cellpose for automated pollen segmentation and quantification of Alexander-stained anther images.

At its core, PAT utilizes Cellpose for instance segmentation and supports both the Cellpose-SAM hybrid (CPSAM) and fined-tuned CPSAM (this study). The application automatically utilizes GPU acceleration when available, with a transparent fallback to the CPU; segmentation of a typical anther image completes in seconds under either configuration. To accommodate varying imaging conditions, PAT offers four channel input modes: full RGB, red channel only, red-minus-green difference, and grayscale. The red-minus-green mode is particularly effective for Alexander-stained material, as it mathematically amplifies the contrast between magenta-stained viable pollen and green-stained background elements. This approach enhances the signal-to-noise ratio and ensures robust performance across varying staining intensities without requiring strict protocol standardization.

A central design principle of PAT is interactive quality control. After automated segmentation, users review results in a synchronized dual-panel display showing the original image alongside a color-coded segmentation overlay. We designed the correction process to be intuitive and rapid: users can remove false detections with a single click or add missed grains by drawing ellipses or freehand polygons. PAT records and propagates all corrections to downstream analyses, creating an auditable record of user interventions.

For laboratories whose imaging conditions differ substantially from our validation set, PAT includes an in-application model training module. Users can fine-tune a new Cellpose model or further train our fine-tuned model directly within the tool using their own corrected images, without writing code or leaving the application. The tool automatically saves the resulting model for future use, enabling progressive adaptation to a laboratory’s specific staining protocols, microscopes, or genotypes.

Once a batch of images is reviewed, PAT performs the statistical analysis automatically. The tool assesses data normality via the Shapiro–Wilk test and subsequently applies either a parametric one-way ANOVA with Tukey’s HSD post-hoc test or a non-parametric Kruskal–Wallis test with Dunn’s post-hoc test (Bonferroni-corrected). Finally, PAT generates a publication-quality figures consisting of a box plot with individual data points, a compact letter display (CLD) for statistical groupings, and significance brackets. Users can export these results as high-resolution PNGs, scalable vector PDFs, or a comprehensive summary report suitable for direct inclusion in manuscripts or laboratory notebooks.

In summary, PAT integrates accurate fine-tuned AI-based segmentation, interactive error correction, laboratory-specific model training, and automated statistical analysis into a single accessible application designed for routine pollen viability quantification.

### 3.4 Application of PAT in discovery of a CAP-D2 allele that suppressed reduced fertility of *smg7-6* mutants

To demonstrate PAT’s utility as a fast and practical tool for genetic discovery, we integrated it into genetic screen to identify novel regulators of meiotic exit. The screen was conducted in the background of *smg7-6* mutant that fails to correctly exit meiosis, leading to a tenfold decrease in viable pollen compared to the wild type and resulting in semi-sterility (Capitao *et al*., 2021). We used PAT to quantify viable pollen from Alexander-stained anthers from pre-selected six candidate lines from B2 population, and *smg7-6* as a control. The application processed the entire dataset and generated publication-quality statistical results in approximately 10 minutes. This high-throughput capability allowed us to objectively rank the strength of pollen viability rescue across all recovered lines with high statistical confidence (**Fig. 4A**).

**Fig. 4.**
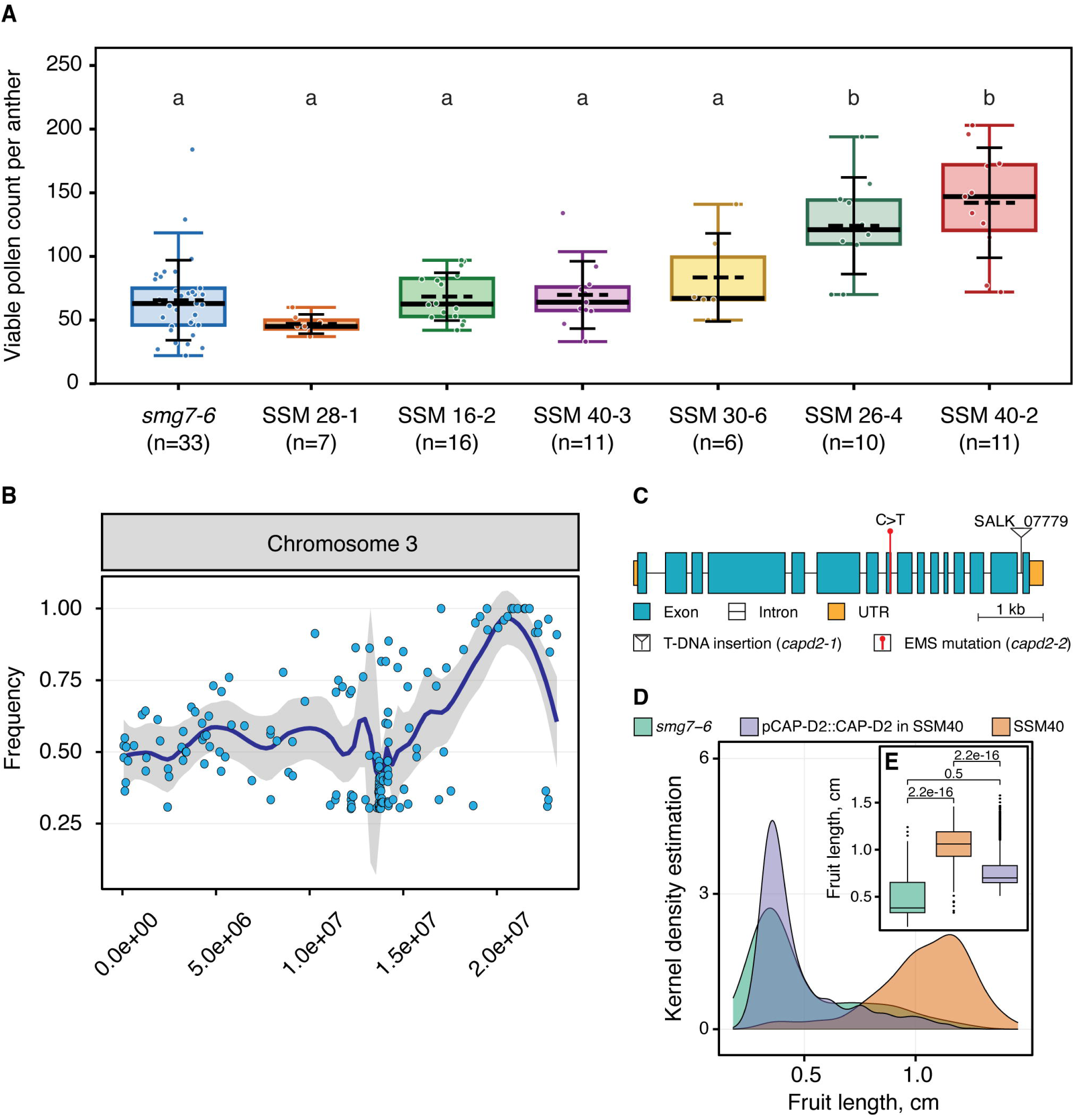
Identification of *CAP-D2* as the suppressor of *smg7-6* via PAT-assisted screening. **(A)** Box plot of pollen quantification across B2 candidate lines using PAT. SSM40-2 shows the most robust rescue of pollen viability, prioritizing it for further characterization. **(B)** Whole-genome SNP association mapping of SSM40-2. Scatter plot represents EMS-induced SNPs on chromosome 3; the y-axis denotes the frequency of the mutant allele in the pooled sequencing data. The purple trend line is calculated using loess regression; gray areas represent 95% confidence intervals. **(C)** Schematic representation of the *CAP-D2* gene structure; the asterisk denotes the position of the p.Ala961Val (C-to-T) mutation. **(D)** Genetic complementation of SSM40-2. Quantification of fruit length across *smg7-6*, SSM40-2, and T1 plants transformed with the *pCAP-D2::CAP-D2* construct. Kernel density estimation and box plots demonstrate that the wild-type transgene reverts the SSM40-2 rescue phenotype to *smg7-6*-like semi-sterility. Box plots show the median and interquartile range (IQR); whiskers extend it 1.5 times IQR. Statistical significance was determined using the Wilcoxon signed-rank test.

The analysis identified line SSM40-2 as having the most robust and statistically significant rescue of pollen counts. Based on this quantitative ranking, SSM40-2 was prioritized for association mapping by whole-genome sequencing (WGS) of B2 segregating population, followed by analysis with artMAP software (Javorka *et al*., 2019). This identified a candidate genomic interval on chromosome 3 (20.5–21.5 Mb) where Ethyl Methanesulfonate-induced SNPs showed strong co-segregation with the suppressor phenotype (**Fig. 4B**). Association mapping in a segregating F2 population further refined this interval to a single SNP in AT3G57060, which encodes the non-SMC HEAT-repeat subunit of the Condensin I complex, CAP-D2 (Schubert *et al*., 2013; Municio *et al*., 2021). This mutation causes an alanine-to-valine substitution (p.Ala961Val) at residue 961 of the protein and exhibited the strongest phenotype-to-genotype association (**Fig. 4C; Fig. S1**).

To confirm that the p.Ala961Val mutation is the causative lesion, we performed a genetic complementation assay. We introduced a wild-type genomic copy of CAP-D2 under its endogenous promoter (*pCAP-D2::CAP-D2*) into SSM40-2 plants. Since the SSM40-2 suppressor allele is recessive, the introduction of a wild-type copy was expected to revert the plants to the *smg7-6* semi-sterile phenotype. As predicted, T1 transformants reverted to semi-sterility; in some instances, plants exhibited complete sterility, likely due to dosage effects from multiple transgene insertions (**Fig. 4D**). This result confirms that substitution of p.Ala961Val in CAP-D2 is the causative mutation in SSM40-2. The allele was outcrossed into wild type background and is hereafter referred to as *capd2-2*.

### 3.5 Mutations in Condensin I, but not Condensin II suppress *smg7-6* meiotic defects

To determine if the suppression is specific to the Condensin I complex, we crossed *smg7-6* with several T-DNA insertion lines. These included an independent *CAP-D2* allele (*SALK_077796*; *capd2-1*) and lines disrupting the Condensin II subunits *CAP-D3* (*SALK_094776*; *capd3*) and *CAP-H2* (*SALK_059304*; *caph2*) (Sakamoto *et al*., 2011; Schubert *et al*., 2013). We again employed PAT to quantify viable pollen from Alexander-stained anthers across all resulting double mutants.

The viable pollen counts revealed a clear specificity: both *capd2-1 smg7-6* and *capd2-2 smg7-6* showed significantly increased pollen viability relative to *smg7-6*, whereas neither Condensin II mutant (*capd3 smg7-6* or *caph2 smg7-6*) exhibited any rescue (**Fig. 5A**). These results demonstrate that suppression is specific to the mutation in Condensin I. Examination of meiotic products in anther lobes further supported this; both *capd2-1 smg7-6* and *capd2-2 smg7-6* produced a significantly higher proportion of tetrads relative to polyads compared with *smg7-6* (**Fig. 5B**). This indicates that a substantial fraction of pollen mother cells (PMCs) in these lines successfully complete meiosis. Notably, the rescue was quantitatively stronger in *capd2-2 smg7-6* than in *capd2-1 smg7-6*.

**Fig. 5.**
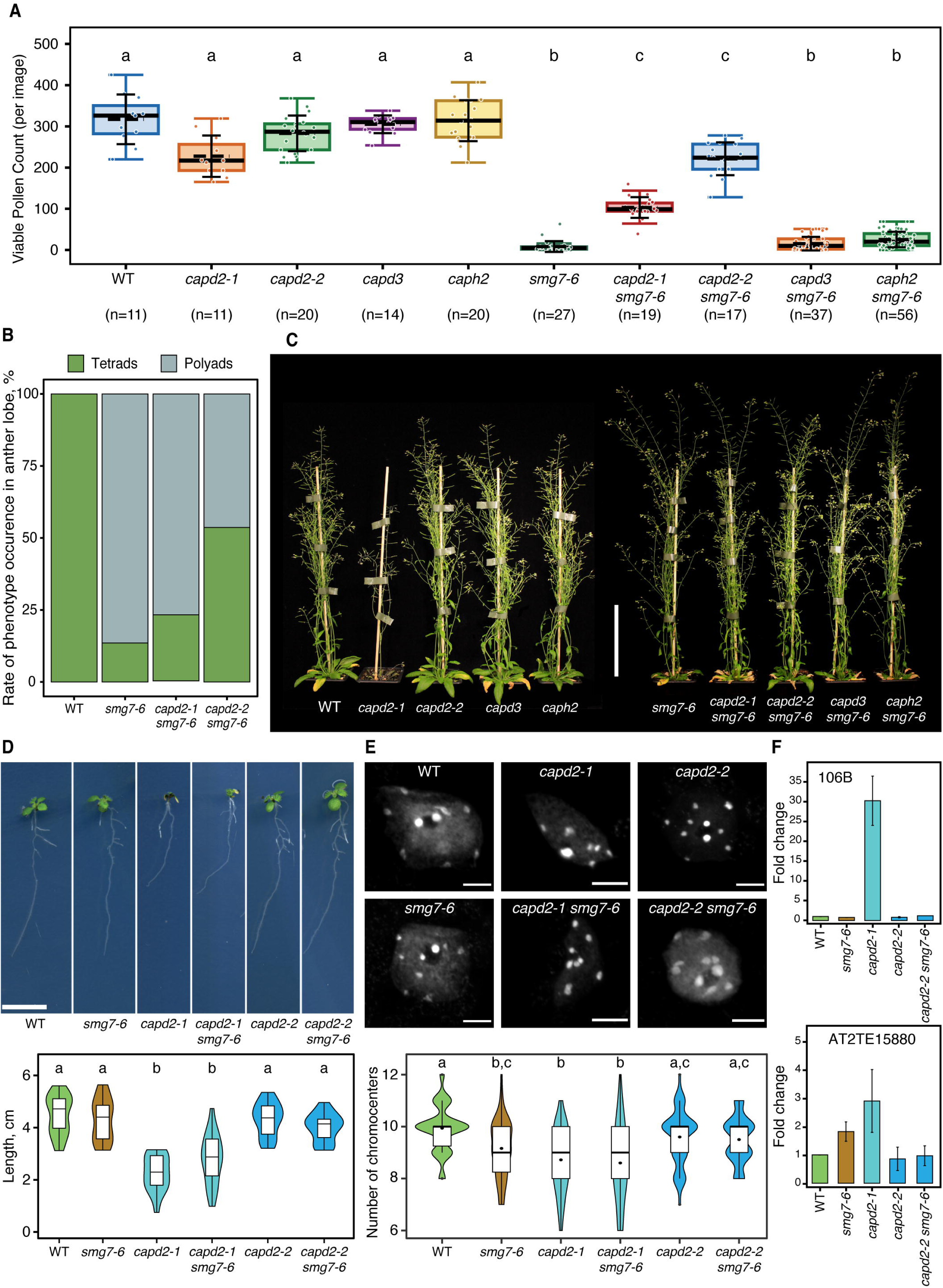
Functional distinction between Condensin I and II in *smg7-6* phenotype suppression. **(A)** Box plot of viable pollen counts across wild-type (WT), Condensin I (*capd2-1*, *capd2-2*), and Condensin II (*capd3*, *caph2*) mutants, both alone and in the *smg7-6* background. **(B)** Quantification of meiotic outcomes (tetrads vs. polyads) in anther lobes (n=20 per genotype), showing reduced polyad frequency in *smg7-6* when combined with *capd2* alleles. **(C)** Phenotype of 45-day-old plants, highlighting the severe vegetative defects of *capd2-1* compared to the normal growth of *capd2-2* and Condensin II mutants. Scale bar = 10 cm. **(D)** Primary root growth analysis showing representative seedling phenotypes (Scale bar = 1 cm) and box plots at 10 days after germination (DAG). Different lowercase letters indicate significant differences (ANOVA, Tukey’s HSD;p < 0.01, n=20). **(E)** Evaluation of interphase chromatin organization. Maximum intensity projections of DAPI-stained nuclei. Scale bar = 5 μm. Quantitative analysis of chromocenter number per nucleus (n=50). Box plots show median and IQR; black points represent group means. Different lowercase letters indicate significant differences (Kruskal-Wallis, Dunn post-hoc; p < 0.01, n=50) (**F**) Quantitative RT-PCR analysis of transposable elements (TEs) *106B* and *AT2TE15880* relative to WT. Error bars represent standard error (SE) of three biological replicates. Data indicate that *capd2-2* retains sufficient Condensin I activity for TE silencing and somatic genome stability.

Since both *capd2-1* and *capd2-2* suppress the *smg7-6* phenotype, we next asked whether they have equivalent effects on somatic growth and Condensin I function. The two alleles diverged sharply in their vegetative development (**Fig. 5C–D**). While *capd2-2* plants remained visually indistinguishable from wild-type throughout their life cycle, *capd2-1* exhibited severe growth defects. At 25 days after germination (DAG), *capd2-1* mutants showed stunted growth and premature bolting compared to wild-type plants, with some individuals failing to survive past this stage (**Fig. S2**). Interestingly, as their above-ground biomass increases, *capd2-1* plants often dislodged from the soil, suggesting insufficient development of root system (**Fig. S3**). Detailed observation in seedlings confirmed that primary root growth in both *capd2-1* and *capd2-1 smg7-6* was affected (**Fig. 5D**). Notably, while *capd2-1 smg7-6* double mutants initially mirrored the stunted phenotype of *capd2-1*, double mutant showed a recovery during the reproductive stage to eventually resemble *smg7-6* single mutants (**Fig. 5C)**. The mechanism underlying this rescue remains unclear. In comparison, the Condensin II subunits *capd3* and *caph2* showed no obvious growth defects (**Fig. 5C**). This comparative phenotyping data suggests that point mutation in *capd2-2* does not fully abolish CAP-D2 function.

To test this at the molecular level, we used chromocenter number and transposable element (TE) expression as readouts for somatic CAP-D2 function (Wang *et al*., 2017). Consistent with previous reports, we found a significant decrease in the number of chromocenters per nucleus in *capd2-1*, reflecting the requirement for Condensin I in heterochromatin compaction (**Fig. 5E**) (Wang *et al*., 2017; Municio *et al*., 2021). Interestingly, while *smg7-6* partially rescued the vegetative defects of *capd2-1*, it failed to rescue these chromocenter defects. In contrast, *capd2-2* and *capd2-2 smg7-6* nuclei showed no major reduction in chromocenter number. Supporting this, qRT-PCR revealed the derepression of Condensin I-regulated TEs (Wang *et al*., 2017) specifically in *capd2-1*, whereas *capd2-2* and *capd2-2 smg7-6* maintained wild-type silencing levels (**Fig. 5F**).

Taken together, these phenotypic and molecular profiles suggest *capd2-2* as a hypomorphic allele. In this mutant, residual CAP-D2 activity is sufficient to maintain essential somatic functions—such as chromocenter integrity, TE silencing, and normal growth yet able to restore meiotic exit in the *smg7-6* background.

## Discussion

This study presents two interconnected contributions to plant reproductive biology. We developed PAT, a graphical desktop application that enables accurate, high-throughput pollen viability quantification without requiring specialist computational infrastructure, and deployed it to identify a hypomorphic allele of CAP-D2 as a suppressor of meiotic exit failure in *smg7-6* (Riehs *et al*., 2008; Schubert *et al*., 2013; Municio *et al*., 2021). Together, these results provide a practical framework for phenotype-driven genetic screening and indicate a role for Condensin I in the final stages of *Arabidopsis* meiosis.

A key insight from our tool development is that general-purpose segmentation models cannot be assumed to transfer reliably to specialized biological imaging tasks. While CPSAM (Marks *et al*., 2025) demonstrated a clear capacity to recognize pollen grains—confirming its underlying architecture is well-suited for this object class—it failed to reliably distinguish pollen from background debris, and non-pollen cells or resolve grains in dense preparations. This was not due to a fundamental limitation of the model, but rather its lack of exposure to the specific colorimetric and morphological context of Alexander-stained anthers. By fine-tuning the model on a curated dataset, we resolved these failure modes. Our results suggest that for most biological applications, fine-tuning an existing foundation model on a modest, domain-specific dataset will consistently outperform the application of a generalist model without adaptation.

PAT specifically addresses the “usability gap” that often prevents high-performing phenotyping tools from seeing broad adoption. While frameworks like Cellpose offer state-of-the-art accuracy, their routine use often requires a level of computational expertise regarding Python environments, library dependencies and command-line workflows that can be a barrier to entry. By distributing PAT as a standalone, cross-platform executable, we have eliminated these technical hurdles. The tool automates the entire pipeline—from batch processing and hardware-accelerated segmentation to statistical analysis and the generation of publication-ready figures with few clicks of mouse. Furthermore, PAT’s integrated retraining workflow ensures the tool remains adaptable; users can “teach” the model to recognize laboratory-specific staining variations without having to annotate datasets from scratch.

Like any computational tool, PAT is not without limitations. In its current implementation, PAT is optimized for TIFF or PNG image formats and assumes the presence of a single anther per image. Users working with proprietary formats such as CZI will therefore need to export their images to a compatible format prior to analysis, and those with multi-anther images should crop them into individual files before batch processing to avoid skewed pollen density calculations. Additionally, the fine-tuned model carries a potential error margin of up to 10 pollen grains, which is particularly relevant for pollen-dense samples such as wild-type anthers. Nevertheless, this limitation is mitigated by an intuitive validation and sanity check module integrated within PAT, enabling users to readily identify and correct any counting discrepancies. Beyond these constraints, PAT is designed with adaptability in mind; its accessible training module and open Cellpose parameters allow users to retrain the model for other cytological staining methods employed in pollen viability analysis, extending its applicability beyond Arabidopsis to potentially other plant species as well.

The utility of PAT was directly demonstrated through its deployment in the EMS suppressor screen in the *smg7-6* background (Capitao *et al*., 2021), where it enabled rapid, objective ranking of pollen viability across the entire cohort in a single session — a task that would otherwise have required days of manual counting and been susceptible to significant observer bias. This quantitative precision was instrumental in prioritizing SSM40-2 for whole-genome sequencing, ultimately leading to the identification of a missense lesion in CAP-D2 (p.Ala961Val), a Condensin I subunit (Schubert *et al*., 2013). The specificity of this suppression to Condensin I is noteworthy. While Condensin II subunits CAP-D3 and CAP-H2 have well-characterized roles in meiotic chromosome organization in Arabidopsis (Hartl *et al*., 2008; Schubert *et al*., 2013; Smith *et al*., 2014; Wang *et al*., 2016), mutations in *capd3* and *caph2* failed to rescue *smg7-6*. Instead, rescue was achieved by two independent alleles of *capd2*, a subunit of Condensin I complex. Condensin I is often associated with heterochromatin compaction, transposable element silencing, and the structural maintenance of centromeres and rDNA (Ono *et al*., 2003; Hirota *et al*., 2004; Smith *et al*., 2014; Wang *et al*., 2017). Whether this reflects a direct role for Condensin I in meiotic exit, or an indirect effect, remains to be determined. Nevertheless, the genetic evidence presented here raises the possibility that Condensin I contributes to the regulation of meiotic progression atleast in *smg7* background.

In conclusion, this work highlights the value of integrating accessible quantitative phenotyping with hypothesis-driven genetic analysis. By removing the counting bottleneck that so often renders large-scale suppressor screens impractical, PAT enabled the objective and efficient identification of *capd2-2* as a suppressor of *smg7-6*, providing a new genetic system with which to dissect the dosage-sensitive requirements of Condensin I across plant development. More broadly, the unexpected link between Condensin I and meiotic exit establishes a new line of investigation into the chromatin-level control of cell cycle transitions in plants, and raises the intriguing possibility that the balance between chromosome condensation and decondensation is a regulated, rate-limiting step in the fidelity of meiotic progression.

## Material and methods

### 1. Plant Material and Growth Conditions

*Arabidopsis thaliana* accession Columbia-0 (Col-0) was used as the wild-type control. The T-DNA insertion lines *smg7-6* (SALK_052532), *capd2-1* (SALK_077796), *capd3-2* (SALK_094776), and *caph2* (SALK_059304) were obtained from the NASC stock center. Seeds were surface-sterilized using chlorine gas for five hours and cultivated on soil (3:1 peat moss to vermiculite) or 0.5× MS medium supplemented with 0.7% agar and 3% sucrose. Plants were grown at 21°C with 50–60% humidity under long-day conditions (16 h light/8 h dark) at a light intensity of 150 μmol m^-2^ s^-1^

### 2. Forward Genetic Screen and Mapping

A forward genetic screen was performed using EMS-mutagenized *smg7-6* seeds to identify meiotic exit suppressors as previously described (Capitao *et al*., 2021; Tanasa *et al*., 2023). Mutants showing improved fertility in the B2 generations were prioritized for whole-genome sequencing (WGS). Genomic DNA was extracted from pooled segregants via the CTAB method, fragmented by sonication to a 100–300 bp range, and prepared for sequencing using the NEBNext® Ultra™ DNA Library Prep Kit. WGS data were analyzed using artMAP to identify candidate SNPs (Javorka *et al*., 2019). The causative mutation was validated through association mapping in a segregating population by correlating phenotype with SNP presence via Sanger sequencing.

### 3. Complementation Assay

For genetic complementation, the genomic sequence of *CAP-D2* (AT3G57060), including 1009 bp of the endogenous promoter and 324 bp of the 3′ UTR, was amplified using Phusion High-Fidelity DNA Polymerase. The fragment was cloned into the pENTR/D-TOPO entry vector and subsequently recombined into the pGWB601 destination vector via Gateway LR Clonase II (Nakagawa *et al*., 2007). The resulting construct was transformed into *Agrobacterium tumefaciens* strain GV3101 and introduced into SSM40-2 plants using the floral dip method (Clough and Bent, 1998). Transgenic T1 plants were selected via BASTA resistance.

### 4. Tetrad Analysis and Microscopy

Meiotic outcomes were evaluated by visualizing callose deposits in whole anthers (Tanasa *et al*., 2023). Inflorescences were fixed in phosphate-buffered saline (PBS) containing 4% formaldehyde under vacuum. Anthers dissected from floral buds smaller than 0.6 mm were stained with a 1:1000 dilution of SCRI Renaissance 2200 (SR2200) in PBS. Anthers were imaged using a Zeiss LSM700 confocal microscope to quantify the frequency of tetrads and polyads.

### 5. Root Length Analysis

To assess somatic growth, seedlings were grown vertically on 0.5× MS plates for 10 days. Primary root growth was monitored by daily scanning of the plates, and root lengths were quantified using ImageJ software (Schneider *et al*., 2012). Statistical significance was determined using ANOVA followed by Tukey’s HSD test (p < 0.01) (Tukey, 1949).

### 6. Alexander Staining

Pollen viability was assessed using Alexander staining to differentiate between viable (magenta) and non-viable (green/blue) grains (Alexander, 1969). Anthers were dissected from large unopened buds and incubated in Alexander staining solution at 50°C for 16–18 hours. Stained anthers were imaged using a ZEISS Axioscope A1 microscope. Viable pollen quantification was performed using the Pollen Analysis Tool (PAT) tool as described in the Results.

### 7. Chromocenter Analysis

Chromocenter numbers were assessed in interphase nuclei isolated from 10-day-old seedlings. Seedlings were fixed in a 3:1 ethanol:acetic acid solution and rehydrated in 100 mM sodium citrate buffer. Following enzymatic digestion with cytohelicase, pectolyase, and cellulase, nuclei were homogenized, sedimented by centrifugation, and fixed onto glass slides with VECTASHIELD containing DAPI. Confocal imaging was performed to quantify the chromocenter frequency per nucleus.

### 8. TE Expression Analysis

Transcript levels of transposable elements (TEs) were analyzed in 7-day-old seedlings. Total RNA was extracted using RNA Blue (Top-Bio), and its quality was verified by MOPS-agarose gel electrophoresis. Following TURBO DNA-free treatment to remove genomic DNA, cDNA was synthesized using SuperScript IV Reverse Transcriptase with oligo(dT) and random hexamer primers. Quantitative RT-PCR was performed on a LightCycler 96 System using KAPA SYBR FAST Master Mix and transcript-specific primers for *106B* and *AT2TE15880* (Wang *et al*., 2017). Expression was normalized to the reference gene *AT4G26410* (Czechowski *et al*., 2005).

### 9. Development of the Pollen Analysis Tool (PAT)

#### Software Architecture and Graphical Interface

PAT was developed as a cross-platform desktop application using Python 3 and the PyQt6 framework. The application architecture utilizes a multi-threaded execution model (via QThread) to decouple the graphical user interface (GUI) from computationally intensive deep-learning inference and image processing tasks, ensuring interface responsiveness during batch operations.

#### Segmentation Engine Instance

The core segmentation engine integrates the Cellpose deep-learning framework (Marks *et al*., 2025), utilizing a U-Net architecture (Ronneberger *et al*., 2015) that predicts per-pixel gradient flows to delineate individual cell boundaries. The tool supports multiple backends, including the generalized the Cellpose-SAM (cpsam) hybrid model and fined tuned cpsam model, which combines a Segment Anything Model (SAM) image encoder with the Cellpose gradient-flow decoder (Stringer *et al*., 2021). For Alexander-stained preparations, a custom channel extraction module was implemented to isolate viable pollen signals, including a “Red-minus-Green” contrast enhancement mode.

#### Image Preprocessing and Algorithmic Filtering

Input images undergo automated normalization using percentile-based contrast stretching (defaulting to the 1st and 99th percentiles) to standardize signal intensity across varying staining batches (Stringer *et al*., 2021). To minimize false positives from non-pollen somatic debris, an automated post-segmentation size filter was implemented. The minimum area threshold is computed dynamically as 0.25 X π X (d/2)^2^ where d is either the user-defined expected grain diameter or the median detected diameter (Auto-detect).

#### Interactive Validation and Model Refinement

An interactive mask correction module was developed using OpenCV and Qt drawing primitives, allowing for the rasterization of elliptical or freehand ROIs directly into the mask array (Bradski, 2000). For model adaptation, a supervised fine-tuning module was integrated using the Cellpose training API (model.trainor train.train_seg), allowing users to perform backpropagation on pre-trained weights using manually corrected ground-truth masks.

#### Batch Processing and Statistical Framework

Parallelization of batch processing is managed via Python’s concurrent.futuresmodule, utilizing a ProcessPoolExecutorwith a “spawn” context to distribute image rendering and segmentation across logical CPU cores. The statistical backend integrates SciPy (Virtanen *et al*., 2020) and statsmodels (Skipper and Josef, 2010) to implement an automated decision tree for comparative analysis. Normality is assessed via the Shapiro–Wilk test (Shapiro and Wilk, 1965); data satisfying parametric assumptions are analyzed via one-way ANOVA followed by Tukey’s HSD (Tukey, 1949), while non-parametric data are analyzed via the Kruskal–Wallis test followed by Dunn’s test with Bonferroni correction (Kruskal and Wallis, 1952; Dunn, 1964). Significant differences are summarized using a Compact Letter Display (CLD) algorithm.

## Acknowledgements/Funding

We acknowledge the the support from CEITEC MU Core facilities Plant Sciences and CELLI of CzechBioimaging supported by MEYS CZ (No. LM2018129). This work was supported by the Czech Science Foundation (22-31712S) and the Ministry of Education, Youth, and Sports of the Czech Republic for the funding from the project TowArds Next GENeration Crops, reg. no.

CZ.02.01.01/00/22_008/0004581 of the ERDF Programme Johannes Amos Comenius.

## Author contributions

DV conducted the EMS screen, identify and characterize the CAP-D2 mutant, contributed in training of the model. VKR conceptualize, design and created PAT. KR supervised the experiment, provided funding and infrastructure necessary for the study. VKR, DV, and KR wrote the manuscript.

## Data availability

Pollen Analysis tool (PAT) is available as open source tool at Github repository (https://github.com/Riha-Lab/Pollen-Analysis-Tool).

## Supplementary Figures

**Fig. S1. EMS mutation in CAP-D2 protein.** (**A**) Schematic representation of CAP-D2 protein structure. Cnd1_N and Cnd1_C are, respectively, conserved N- and C-terminal domains of condensin subunit I. (**B**) Protein alignment of CAP-D2 across the tree of life. Red box highlights the position corresponding to Arabidopsis alanine 961 (Ala961).

**Fig. S2.** A representative picture showing phenotypes of 25-days-old wild type (WT), *smg7-6*, *capd2-1, capd2-1 smg7-6, capd2-2* and *capd2-2 smg7-6*. Scale bar = 5 cm.

**Fig. S3.** A representative picture showing roots of *capd2-1* plant after dislodgement from the pot.

